# HY5 is not integral to light mediated stomatal development in Arabidopsis

**DOI:** 10.1101/756221

**Authors:** Nicholas Zoulias, Jordan Brown, James Rowe, Stuart A. Casson

**Author notes:** Chameleon Communications International, 50 Alderley Rd, Wilmslow, UK.

## Abstract

Light is a crucial signal that regulates many aspects of plant physiology and growth including the development of stomata, the pores in the epidermal surface of the leaf. Light signals positively regulate stomatal development leading to changes in stomatal density and stomatal index (SI; the proportion of cells in the epidermis that are stomata). Both phytochrome and cryptochrome photoreceptors are required to regulate stomatal development in response to light. The transcription factor *ELONGATED HYPOCOTYL 5* (*HY5*) is a key regulator of light signalling, acting downstream of photoreceptors. We hypothesised that HY5 could regulate stomatal development in response to light signals due to the putative presence of HY5 binding sites in the promoter of the *STOMAGEN* (*STOM*) gene, which encodes a peptide regulator of stomatal development. Our analysis shows that HY5 does have the potential to regulate the *STOM* promoter *in vitro* and that HY5 is expressed in both the epidermis and mesophyll. However, analysis of *hy5* and *hy5 hyh* double mutants (*HYH*; *HY5-HOMOLOG*), found that they had normal stomatal development under different light conditions and the expression of stomatal developmental genes was not perturbed following light shift experiments. Analysis of stable lines overexpressing HY5 also showed no change in stomatal development or the expression of stomatal developmental genes. We therefore conclude that whilst HY5 has the potential to regulate the expression of *STOM*, it does not have a major role in regulating stomatal development in response to light signals.

## Introduction

Stomata are the microscopic pores on the epidermal surface of leaves and they are vital for regulating plant gas exchange, which is achieved via regulation of the stomatal pore aperture in response to changes in the local environment (reviewed in Assmann and Jegla, 2016 (1)). Our understanding of stomatal development has advanced significantly over recent years, particular in the model dicot, *Arabidopsis thaliana* (reviewed in Zoulias et al. 2018 (2)). The basic helix-loop-helix (bHLH) transcription factors SPCH, MUTE and FAMA positively regulate three formative steps in stomatal development. SPCH regulates entry into the lineage, MUTE is required to produce the immediate precursor of guard cells and FAMA regulates guard cell formation (3–5). These transcription factors form dimers with the bHLH proteins ICE1 and SCRM2 to regulate target genes (6). A ligand/receptor/MAP kinase pathway negatively regulates stomatal development by targeting the SPCH-ICE1 dimer (7, 8). Stomatal lineage cells produce secreted peptides (EPF1 & EPF2), which bind a receptor complex that includes members of the ERECTA family (ERf), the TOO MANY MOUTHS (TMM) receptor-like protein and members of the SERK family of receptor kinases (9, 10). This activates a MAP kinase pathway that phosphorylates SPCH, targeting it for degradation (7, 8). STOM belongs to the same family as EPF1/2 however, it positively regulates stomatal development by competing with EPF1/2 to bind the receptor and switch off the MAPK pathway (9, 11). Unlike EPF1 and EPF2, STOM is not expressed in the epidermis but is secreted from the mesophyll.

The number of stomata that develop on new leaves is regulated by environmental conditions, and light quantity and quality have been shown to mediate changes in stomatal development via the red/far-red light perceiving phytochromes and the blue light perceiving cryptochromes (12, 13). For example, light quantity positively regulates stomatal development leading to an increase in stomatal index. Whereas, *phyB* mutants are defective in mediating responses to light quantity and have a significantly reduced stomatal index compared to wild type plants (12). Photoreceptors regulate the activity of COP1, an E3 ubiquitin ligase, that targets positive regulators of photomorphogenesis for degradation such as the transcription factor HY5 (14–16). HY5 is a transcriptional activator and repressor and has been shown to be a key regulator of light signalling (17), sometimes acting redundantly with the closely related transcription factor, HYH (16). An analysis of HY5 binding sites determined that there were >3000 binding sites within the Arabidopsis genome (18). An analysis of this data revealed that HY5 binds to the *STOM* promoter presenting an attractive hypothesis that HY5 may regulate stomatal development by regulating expression of *STOM*. Here, we have used a combination of genetic and molecular analyses to investigate any potential role of HY5 in regulating stomatal development in response to light signals. However, whilst our data indicates that HY5 has the potential to regulate *STOM* expression *in vitro*, it does not appear to play a major role in light regulation of stomatal development.

## Results

An analysis of HY5 ChIP data (18) revealed that HY5 has potential binding sites within ~250bp of the transcriptional start site (TSS) of the *STOM* gene (TSS chm 4, 7587099 bp; oligo sites 7586680, 7587343). Z-boxes are light responsive elements found in a number of light responsive genes and have been implicated as HY5 binding sites (29). Analysis of the *STOM* promoter identified two putative Z-boxes (-151 and -601 from the ATG) as well as a GATA-box (15) (- 126 from the ATG) that could account for the HY5 ChIP result (Figure 1A). To test whether HY5 has the potential to directly regulate the expression of *STOM* we first used a dual luciferase assay (see methods) and co-transformed protoplasts with *CaMV35SproHY5* and *STOMpromLUC* constructs. As a positive control, we also co-transformed protoplasts with *CaMV35SproHY5* and *HY5promLUC* constructs, as HY5 has been shown to binds its’ own promoter (ACE-box -282 from the ATG) to autoregulate expression, which was confirmed in these experiments (Figure 1B; (30)). In this i*n vitro* assay, HY5 can clearly regulate *STOM* expression supporting the ChIP data (Figure 1B).

**Fig 1.**
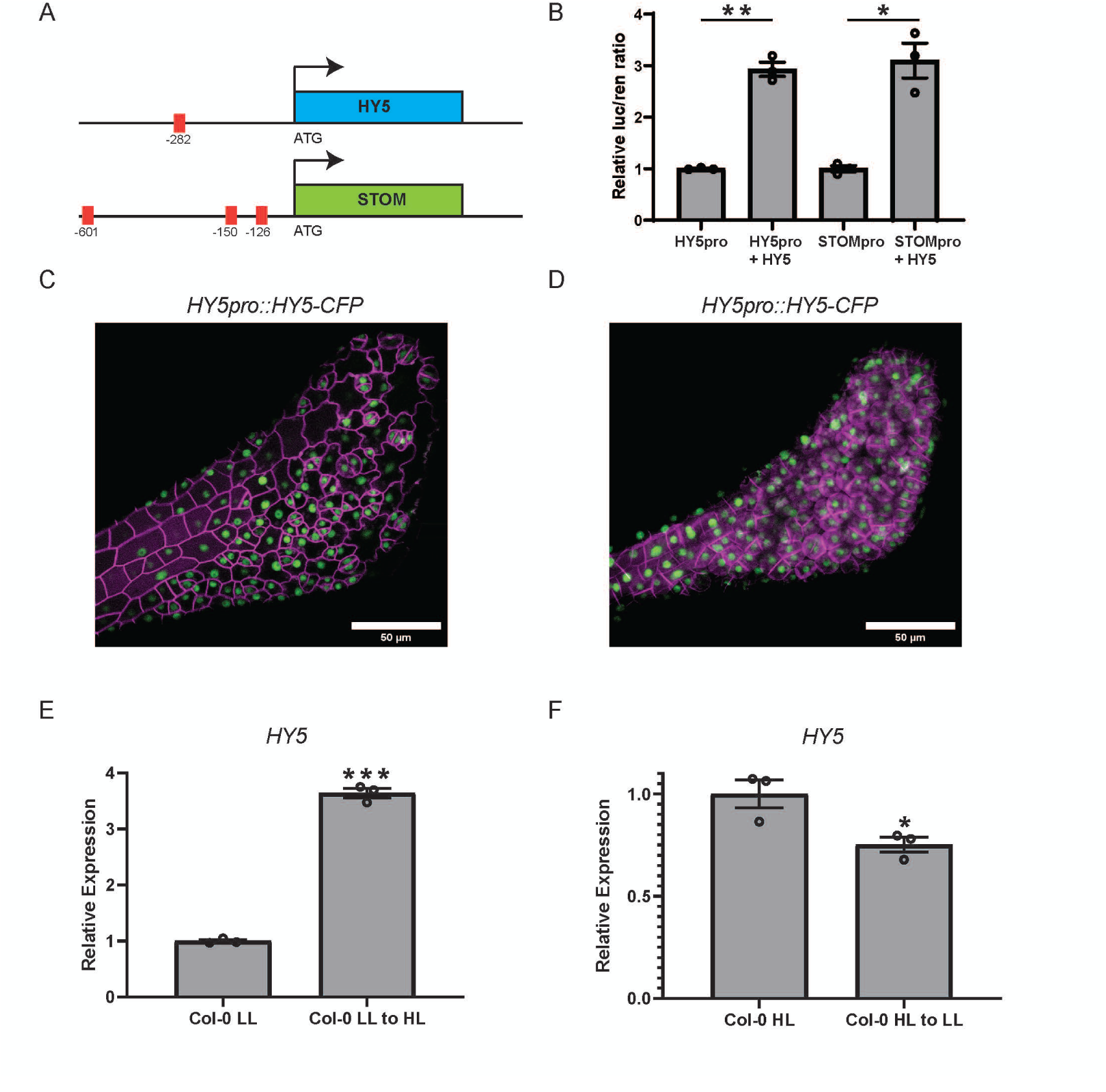
HY5 expression and regulation in response to light. (A) Diagrammatic scheme of *STOMAGEN* and *HY5* promotors with putative HY5 binding boxes highlighted in red. *HY5* promotor, ACE-box -282 bp from the ATG. *STOMAGEN* promotor, Z-boxes -151 and -601 as well as a GATA-box – 126 bp from the ATG. (B) Dual luciferase assays showing relative expression of *HY5pro::Luc 35Spro:: Renilla* and *STOMpro::Luc 35Spro::Renilla* transfected into *Arabidopsis* protoplasts with and without *35Spro::HY5*. All experiments were completed with biological and technical triplicate, error bars indicate SEM. Asterisks indicate significant differences between promotors only and promotors plus *35S::HY5*. (Welch’s t-test: *P < 0.05; **P < 0.01). (C) Confocal Z-sum projection of *HY5pro::HY5-CFP* (green) expression in wave_138Y (Col-0, magenta) in the epidermis (digitally segmented). (D) Confocal Z-sum projection of *HY5pro::HY5-CFP* (green) expression in wave_138Y (Col-0, magenta) in the mesophyll (digitally segmented). (E) Relative expression levels of *HY5* in low light (50 μmol m^−2^ s^−1^) and 6 hours after a low to high light transfer (50 to 250 μmol m^−2^ s^−1^) examined by qRT-PCR, *UBC21* served as internal control. Experiments were performed in biological triplicate and error bars indicate SEM. Asterisks indicate significant differences between low light and low to high light transfers. (Welch’s t-test: ***P < 0.001). (F) Relative expression levels of *HY5* in high light (250 μmol m^−2^ s^−1^) and 6 hours after a high to low light transfer (250 to 50 μmol m^−2^ s^−1^) examined by qRT-PCR, *UBC21* served as internal control. Experiments were performed in biological triplicate and error bars indicate SEM. Asterisks indicate significant differences between low light and low to high light transfers. (Welch’s t-test: *P < 0.05).

Given that HY5 has the potential to regulate *STOM* expression we next examined expression of HY5 using confocal microscopy to study the localisation of HY5:CFP in HY5proHY5:CFP transgenic plants. It should be noted that previous studies have shown that HY5 is expressed widely throughout the plant including leaf tissue and that HY5 protein is mobile, therefore this already indicates that HY5 has the potential to regulate STOM expression in the mesophyll (31, 32). Our analysis confirmed the results of previous studies and HY5:CFP signal was clearly detected in the epidermis and stomatal lineage cells as well as the mesophyll (Figure 1C,D). Therefore, HY5 has the potential to regulate *STOM* expression as determined by dual luciferase experiments and is expressed in the relevant tissues.

We next examined whether HY5 is required for light regulation of stomatal development, first examining the transcriptional response of *HY5* to changes in irradiance. Seedlings were grown under low (LL; 50 µmol m^−2^ s^−1^) or high (HL; 250 µmol m^−2^ s^−1^) before transferring to the alternate irradiance for 6h to investigate LL-HL and HL-LL responses. These conditions were chosen because plants grown under these steady state conditions have significant differences in their stomatal development (see Figure 2A, B). *HY5* expression was significantly affected by changes in irradiance with significant upregulation following a LL-HL transfer and downregulation following a HL-LL transfer (Figure 1 E, F). Under these same experimental conditions, we see a similar trend for the major regulators of stomatal development *SPCH* and *MUTE* as well as *STOM* (Figure 2C, D). Therefore, *HY5* and these stomatal regulators show dynamic changes in expression following light shifts, which correlate well with the positive impact of light on stomatal development.

**Fig 2.**
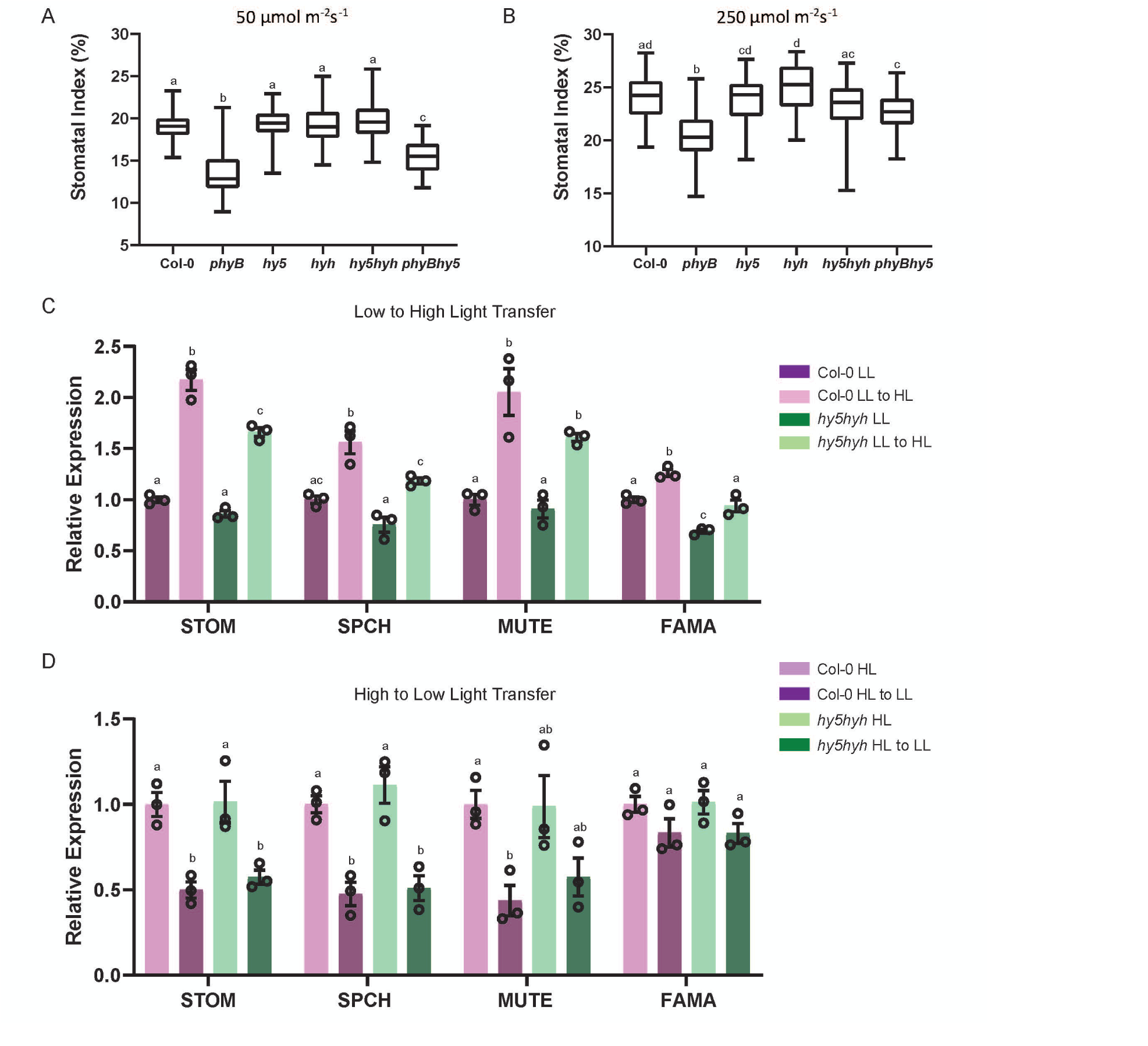
HY5 is not involved in light regulation of stomatal development. (A) Stomatal index of Col-0, *phyB*, *hy5*, *hyh*, *hy5hyh*, and *hy5phyB* mature leaves grown under low light (50 µmol m^−2^ s^−1^). Box plots indicate the 25^th^ and 75^th^ quartiles with the line representing the median; whiskers are the minimum and maximum range. One-way ANOVA was performed to test statistical difference; letters denote significance with a posthoc Tukey test. Alpha = 0.05. (B) Stomatal index of Col-0, *phyB*, *hy5*, *hyh*, *hy5hyh*, and *hy5phyB* mature leaves grown under high light (250 µmol m^−2^ s^−1^). Box plots indicate the 25^th^ and 75^th^ quartiles with the line representing the median; whiskers are the minimum and maximum range. One-way ANOVA was performed to test statistical difference; letters denote significance with a posthoc Tukey test. Alpha = 0.05. (C) qRT-PCR relative expression levels of transcription factor regulators of stomatal development (*STOM, SPCH*, *MUTE*, *FAMA*) in Col-0 and *hy5hyh* backgrounds following a transfer from low to high light (50 to 250 μmol m^−2^ s^−1^). Experiments were performed in biological triplicate and error bars indicate SEM. One-way ANOVA was performed to test statistical difference for each gene tested; letters denote significance with a posthoc Tukey test. Alpha = 0.05. (D) qRT-PCR relative expression levels of transcription factor regulators of stomatal development (*STOM, SPCH*, *MUTE*, *FAMA*) in Col-0 and *hy5hyh* backgrounds following a transfer from high to low light (250 to 50 μmol m^−2^ s^−1^). Experiments were performed in biological triplicate and error bars indicate SEM. One-way ANOVA was performed to test statistical difference for each gene tested; letters denote significance with a posthoc Tukey test. Alpha = 0.05

To further examine the role of HY5 in light regulation of stomatal development, we next performed epidermal cell counts on plants grown under our LL and HL conditions. For these analyses, we examined *hy5* mutants as well as a *hyh* and a *hy5hyh* double mutant, in case of any functional redundancy between these closely related transcription factors. RT-PCR analysis of the *hy5 hyh* double mutant revealed that it is null for both of these genes, whilst hypocotyl measurements clearly show the redundant role both these genes play in hypocotyl elongation (Figure S1A-C). *phyB-9* mutants were also included in these analyses as phyB has been demonstrated to be required for light mediated stomatal development, with reduced stomatal densities and stomatal index when grown at HL (Casson et al. 2009). Stomatal index measurements indicate whether a factor influences decisions in the stomatal developmental pathway and under both our LL and HL conditions, none of *hy5*, *hyh* and *hy5hyh* showed any difference to the Col-0 control, whereas *phyB* mutants had significantly reduced SI (Figure 2A, B). *hy5* mutants also did not have any impact on stomatal density in these conditions (Figure S2A, B). Analysis of a *phyBhy5* double mutant showed a phenotype similar to that of the *phyB* single mutant (Figure 2A, B).

These analyses were performed on mature leaves so we also examined stomatal development in cotyledons to see if HY5 or HYH might influence stomatal development in these early organs. However, epidermal counts of the *hy5hyh* mutant revealed no difference to the WT (Figure S2 C). Therefore, phenotypic data from both mature and juvenile leaf material suggests that HY5 (and HYH) are not likely to have a significant role in regulating stomatal development under these conditions.

Given the fact that HY5 has the potential to regulate *STOM in vitro* but does not appear to have a major role based on phenotypic analysis, we used qRT-PCR to investigate whether HY5 regulates changes in the expression of *STOM* following changes in irradiance. For these analyses, the *hy5hyh* double mutant was compared to the Col-0 control and both LL-HL and HL-LL transfers were examined (Figure 2C, D). In line with the stomatal counts data, our gene expression analysis revealed that with regards the genes analysed, including *STOM*, the *hy5hyh* double mutant responded as the control indicating that these factors are not required for the dynamic light mediated changes in expression of these key stomatal developmental genes. To investigate further, we next examined lines that stably overexpress *HY5* (*35Spro::HY5*). The *35Spro::HY5* lines showed high level expression of *HY5* under both LL and HL conditions (Figure S2E, F). If HY5 regulates *STOM* expression *in planta* we therefore would predict changes in *STOM* expression in these lines, however qRT-PCR analysis did not support this, again indicating that HY5 is unlikely to have a major role regulating *STOM* expression *in planta* (Figure S2E, F). This is further supported by phenotypic data of the *35Spro::HY5* which shows no change in SI when compared to Col-0 (Figure S2D).

## Discussion

Stomatal development is under environmental control and previous studies have demonstrated that phytochrome and cryptochrome photoreceptors as well as the negative regulator of photomorphogenesis, COP1, are required for light mediated control (12, 13). HY5 is an integral regulator of transcriptional responses to light signals and functions downstream of the photoreceptors and is targeted directly, along with HYH, by COP1 (14–17). A genome wide analysis of HY5 binding sites indicated that HY5 has the potential to bind to the *STOM* promoter (18). As STOM is a positive regulator of stomatal development, this presents an attractive hypothesis that light signals regulate stomatal development via HY5 regulation of *STOM*. As COP1 activity is mediated by the photoreceptors then it would be expected that *STOM* expression would be upregulated under higher irradiances and reduced in lower light conditions. Indeed, our dynamic light transfer experiments clearly show that *STOM* expression is regulated in this manner. Using *in vitro* methods we can also demonstrate that HY5 has the potential to positively regulate *STOM*. However, systematic testing of this hypothesis using a range of molecular and genetic tools indicates that HY5, as well as HYH, are not major regulators of stomatal development. If this were the case, then we would expect changes in stomatal development in single or double *hy5* and *hyh* mutants. In conditions where a *phyB* mutant clearly shows reduced SI, as expected from previous studies, *hy5*, *hyh* and *hy5hyh* mutants have WT-like phenotypes despite clearly showing the expected elongated hypocotyl phenotype previously shown for these mutants. Furthermore, if HY5 was integral for *STOM* regulation *in planta*, we would again expect *hy5* mutants or overexpressing lines to show defective regulation of *STOM* in response to light signals but this is not the observation.

How then can we account for the ChIP data and lack of HY5 mediated responses *in planta* in these studies? It is possible that caution should be applied to the ChIP data as this study was performed with a line overexpressing HY5 and hence may not reflect native HY5 binding sites. Alternatively, HY5 may indeed bind the *STOM* promoter but its action is inhibited by other factors; for example MONOPTEROS/ARF5 are known to negatively regulate *STOM* (33). However, given that *phyB* mutants are defective and *hy5* mutants are not, it would indicate that HY5 is not an integral component of light regulated stomatal development. The recent studies that have shown that ICE1 is a target of COP1 and ICE1’s pivotal role in regulating SPCH could account for some aspects of light mediated stomatal development (8, 34).

## Materials and Methods

### Plant materials and growth conditions

The Arabidopsis ecotype Col-0 was used as the wild-type control in all experiments. The following mutants and transgenic lines were used in this study; *phyB-9* (19), *hy5* (salk_096651, (20)), *hyh* (WiscDsLox253D10), *HY5proHY5:CFP* and *35S:HY5*. The *hy5 hyh* double and *phyB hy5* mutants were generated by crossing the respective lines and double mutants selected in the F2 generation. DNA was extracted (21) and the genotypes verified using the primers Salk Lba1, hy5salk_096651cFor and hy5salk_096651cRev for *hyh* and LBp745DsLox, HYHwiscFor and HYHwiscRev for *hyh*. The *phyB-9* point mutation was verified by sequencing of a PCR product generated using the primers phyBWTfor and phyBWTrev.

For stomatal counts and qRT-PCR analysis, plants were grown on Levingtons F2+sand compost in environmental control chambers (Conviron BDR16) fitted with 22x Philips Master Pl-L 55W/84°/4P bulbs at an irradiance of 50 to 250 μmol m^−2^ s^−1^, a 11 h photoperiod (20°C day and 16°C night) and 65% RH. All light transfers were performed two hours post dawn. For low to high light transfers, seedlings (8 days post germination) were moved from an irradiance of 50 to 250 μmol m^−2^ s^−1^ for six hours. For high to low light, transfers seedlings (10 days post germination) were moved from an irradiance of 250 to 50 μmol m^−2^ s^−1^ for six hours.

For analysis of hypocotyl lengths, Col-0, *hy5*, *hyh*, and *hy5hyh* were grown on ½ strength Murashige and Skoog medium (0.8% plant agar) for 6 days post germination in environmental control chambers (Conviron BDR16) fitted with 22x Philips Master Pl-L 55W/84°/4P bulbs at an irradiance of 150 μmol m^−2^ s^−1^, a 11 h photoperiod (20°C day and 16°C night) and 65% RH. For confocal imaging, *HY5proHY5:CFP* seedlings were grown on ½ strength Murashige and Skoog medium (0.8% plant agar) for 5 days post germination in environmental control chambers (Conviron BDR16) fitted with 22x Philips Master Pl-L 55W/84°/4P bulbs at an irradiance of 150 μmol m^−2^ s^−1^, a 11 h photoperiod (20°C day and 16°C night) and 65% RH.

### HY5proHY5:CFP construction

HY5 genomic sequence was amplified from Col-0 gDNA using using Q5® High-Fidelity DNA Polymerase (New England Biolabs) and the primers HY5proFor-Not and HY5Rev-Sal. The PCR fragment was digested with *Not*I and *SalI* and ligated into *NotI*-*SalI* digested vector pGKGWC (22). This binary vector was co-transformed with the helper plasmid pSOUP into Agrobacterium C58C1 before transformation of the wave_138Y plasma membrane marker line (Columbia background;(23)) using the floral-dip method (24).

### Confocal Imaging

Seedlings were mounted in ddH_2_O without the hypocotyl and root. The young developing leaves were imaged with an Olympus FV1000 confocal laser scanning microscope with a 60X oil lens, producing Z-stacks through the abaxial epidermis and into the mesophyll layers. CFP was excited with the 440 nm laser line. YFP was excited with the 515 nm laser line. Microscope settings were not changed between cotyledons of the same line to ensure cross comparability. Images shown are representative of N = 8 images.

### Image Rendering

A plugin (EZ-Peeler v 0.16) was written for ImageJ to segment plant surface and to extract the data from a user defined depth and offset below this contoured surface. This allows separate rendering of epidermis and mesophyll, confocal images are Z sum projections of these segmented images. HY5-CFP channel intensity is consistent between images, wave_138Y brightness has been optimised in each image to cell outlines to be easily visible and not obscure the HY5-CFP channel. EZ-Peeler was developed in Python for ImageJ. Source code is available at https://github.com/JimageJ/EZ-Peeler.

### CaMv35S:HY5 (35Spro::HY5) construction

The *HY5* CDS was amplified from Col-0 cDNA using HY5-AscI-For and HY5-PacI-Rev. The amplified *HY5* fragments and pMDC32 (25) were both digested with PacI and AscI and ligated together. This binary vector was transformed into Agrobacterium C58C1 before transformation of Col-0 plants using the floral dip method (24).

### Stomatal Impressions and counting

For mature leaf counts, 15 fully expanded and healthy mature leaves (three leaves from five plants, per plant line) were selected for cell counts. Dental resin (coltene, PRESIDENT, light body dental resin) was applied to the abaxial surface of the leaves and allowed to set. Leaf material was removed and impressions coated with one layer of clear nail varnish. Clear tape was placed over the clear nail varnish and mounted on to slides for microscopic imaging. A Leica DM IRBE Inverted Microscope with Planachromat 20x/ 0.4∞/ 0.17-A lens was used to image impressions. Micro-Manager 1.4 software was used to acquire Z-stack files of 3 points on a mature leaf (base (b), middle (m) and tip (t)). Each Z-stack file was opened through ImageJ software, and a 400µm × 400 µm region of interest chosen for counting.

Impressions of the abaxial surface of cotyledons were made using dental resin (Impress Plus Wash Light Body, Perfection Plus Ltd, Totton, UK). Clear nail varnish was applied to the set impression after removal from the cotyledon, and Z-stack images captured at 20X on a Brunel n300-M microscope equipped with a Prior ES10ZE Focus Controller and Moticam 5 camera. 30 cotyledons for each genotype (area 0.24 mm-2) were examined per experiment and statistical analysis performed using GraphPad Prism.

Stomatal Index (SI) = (total stomata/ total stomata+ total epidermal cells) X 100

Stomatal Density (SD) = total stomata/mm^2^

### Hypocotyl Measurements

Hypotcotyls were imaged using a Lecia S9i stereo microscope with an integrated camera. ImageJ software was used to measure hypocotyl length. Experiments were repeated in triplicate (N ≥ 101) and statistical analysis performed using GraphPad Prism.

### RNA extraction, RT-PCR and qRT-PCR

For all RT-PCR and qRT-PCR experiments, roughly 100mg of seedling or leaf tissue (approximately 20 seedlings) was collected in 2 ml safe lock tubes containing a 5 mm steel ball bearing and flash frozen in liquid nitrogen. Frozen tissue was disrupted in a TissueLyser II (Qiagen; Manchester, UK) and RNA extracted using a Quick-RNA™ MiniPrep kit (#R1055a; Zymo Research, Irvine, USA) according to the manufacturer’s instructions including an on column DNase step. 2μg of total RNA was reverse transcribed using the High-Capacity cDNA Reverse Transcription kit (#4368814; Applied Biosystems, Foster City, USA). cDNA was diluted 20X in ddH2O prior to RT-PCR or qPCR. Q5® High-Fidelity DNA Polymerase (New England Biolabs) was used for RT-PCR, performed with a T100 thermal cycler (Bio-Rad, Watford, UK) with 35 cycles of 95°C-20s, 57°C-20s 72°C-30s. RT-PCR results were visualised using a 1% agarose gel with ethidium bromide (0.5 μg/mL). SYBR® Green JumpStart™ Taq ReadyMix (#S5193; Sigma-Aldrich, Poole, UK) was used for qRT-PCR (3.5mM MgCl2; 375nM primer, see Table S1 for primer sequences) and was performed using a CFX Connect Real-Time PCR Detection System (Bio-Rad, Watford, UK) with 40 cycles of 95°C-10s, 57°C-10s 72°C-15s and a final dissociation curve. Relative expression of target genes in the different samples was calculated from UBC21 or UBQ10 normalized target signals using the ΔΔCT method (26), statistical analysis performed using GraphPad Prism.

### Dual luciferase

#### Plasmid construction

*HY5proLUC* and *STOMproLUC* in pGREEN-800 Luc (27) were created using standard cloning protocols. In brief, a fragment ~2kbp upstream of the ATG was cloned from Col-0 genomic DNA using Q5 polylmerase and specific primers (see Table S1 for all primers sequences). The amplified promotor fragments and pGREEN-800 Luc were both digested with KpnI and NcoI and ligated. For *35SproHY5* in pDH51-YFPc (22), *HY5* CDS was amplified from Col-0 cDNA using Q5 and the specific primers HY5BamHI-For and HY5XhoI-Rev. Both the *HY5* CDS fragment and pDH51-YFPc were digested with BamHI and XhoI prior to ligation.

#### Protoplast Isolation

Protoplasts were isolated from mature leaves (7-10 leaves per isolation), of 4-5 week old Col-0 plants grown in an irradiance of 250 μmol m^−2^·s^−1^, using the ‘Tape-*Arabidopsis*-Sandwich’ method (28). Autoclave tape was affixed to the adaxial surface and the excess tape carefully removed. Another strip of autoclave tape was affixed to the abaxial surface and then carefully peeled away, removing the epidermis and exposing the mesophyll layers. Peeled leaves were incubated in 10mL of enzyme solution [1% cellulase ‘Onozuka’ R10 (Duchefa Biochemie, Netherlands), 0.25% macerozyme ‘Onozuka’ R10 (Duchefa Biochemie, Netherlands), 0.4 M mannitol, 10 mM CaCl_2_, 20 mM KCl, 0.1% BSA and 20 mM MES, pH 5.7] for one hour on a shaking platform (50 rpm). After one hour the solution now containing protoplasts was removed and centrifuged for three minutes at 100 × *g*, and then washed twice with 25 mL of W5 buffer (154 mM NaCl, 125 mM CaCl_2_, 5 mM KCl, 5 mM glucose, and 2 mM MES, pH 5.7). Protoplasts were counted using a hemocytometer and following a centrifugation at 100 × *g* (one minute) were resuspended in MMg solution (0.4 M mannitol, 15 mM MgCl2, and 4 mM MES, pH 5.7) to a final cell density of 5×10^5^ cell/mL. 10-20µg of plasmid was mixed with 1 × 10^5^ protoplasts in MMg solution at room temperature. An equal volume of freshly prepared PEG solution (40% PEG MW 4000, 0.1 M CaCl2 and 0.2 M mannitol) was added to the protoplast/plasmid mixture and allowed to incubate for 10 minutes at room temperature. Following the incubation, the protoplasts were wash three times with W5 buffer and centrifuged at 100 × *g* for one minute in between washes. After the final wash the protoplasts were resuspended in one mL of W5 buffer and incubated in the original growth conditions for 16-24 hours.

#### Dual Luciferase Assays

Protoplasts that had been transfected with 10 µg of each individual construct or equivalent volume of water for the negative controls were used for the dual luciferase assay. 16-24 hours after transfection, protoplasts were pelleted by centrifugation (14,000 × *g* for 30 seconds). Protoplasts were lysed in 1x passive lysis buffer (the Dual-Luciferase® Reporter Assay System, Promega) and incubated at room temperature for 15 minutes. Following the incubation, 20µL (approximately 6.6x 10^4^ cells) of lysed protoplasts were added to 100 µL LARII (the Dual-Luciferase® Reporter Assay System, Promega) vortexed briefly and measured immediately in a luminometer (Sirius Luminometer, Berthold Detection Systems). The luminescence of Luciferase was quenched and Renilla luminescence measured by the addition of 100 µL of Stop & Glo® Buffer (the Dual-Luciferase® Reporter Assay System, Promega). Luminescence was measured in technical triplicates for all combinations of transfected plasmids and each transfection was repeated thrice, statistical analysis performed using GraphPad Prism.

## Acknowledgements

We would like to thank Roger Hellens for kindly providing materials. This work was funded by the Biotechnology and Biological Sciences Research Council (BBSRC) grant (BB/N002393/1) and a Departmental PhD studentship (JB) to SAC.

## Supporting Information

**S1 Fig.**
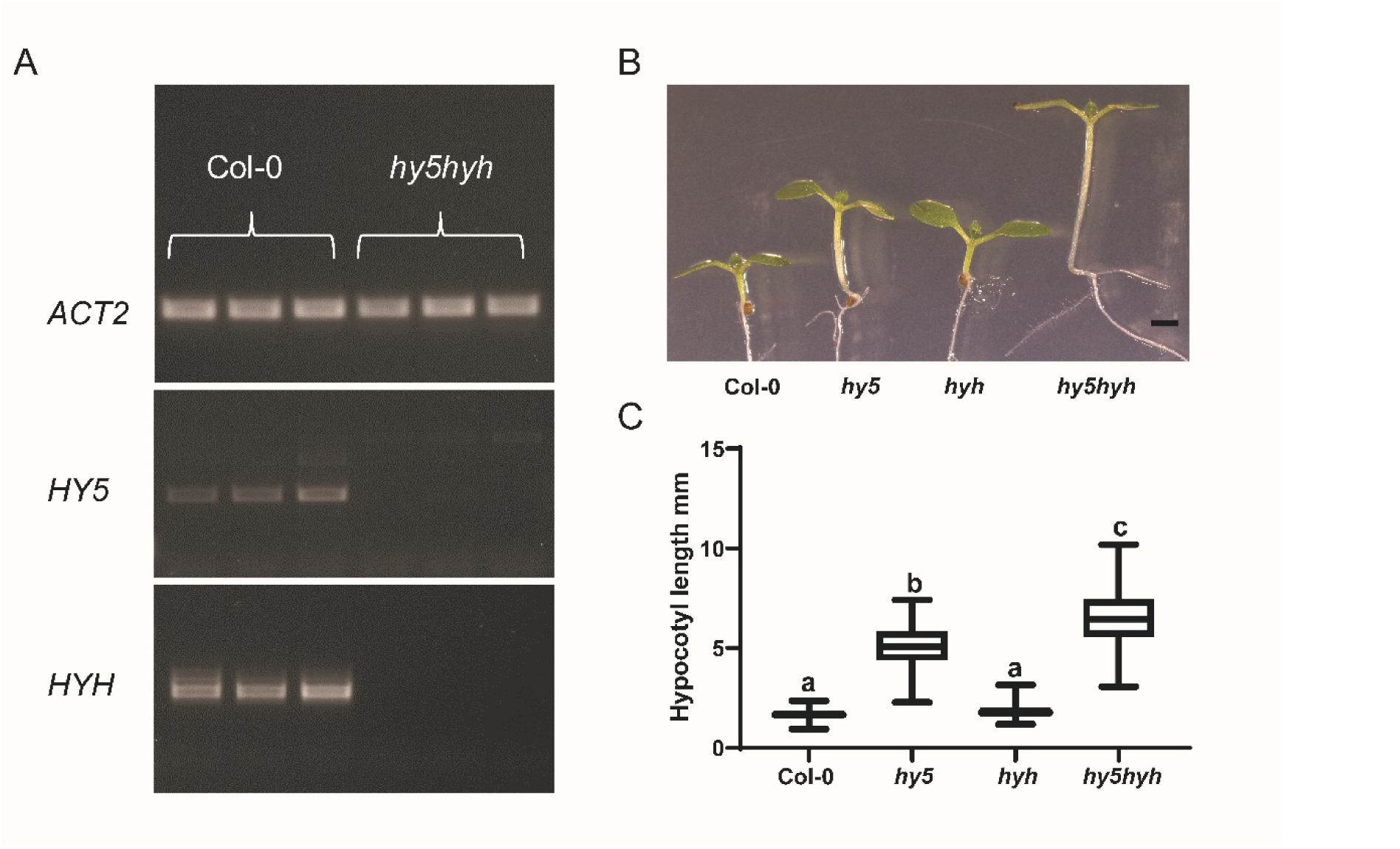
Genotyping and phenotyping the *hy5hyh* mutant. (A) RT-PCR of full length *HY5* and *HYH* in the Col-0 and *hy5hyh* backgrounds. Three biologically independent samples were used. (B) Representative image of Col-0, *hy5*, *hyh* and *hy5hyh* hypocotyls. Scale bar is 1.5 mm in length. (C) Hypocotyl length measurements of Col-0, *hy5*, *hyh* and *hy5hyh* grown in 150 μmol m^−2^ s^−1^ of light. Box plots indicate the 25^th^ and 75^th^ quartiles with the line representing the median; whiskers are the minimum and maximum range. One-way ANOVA was performed to test statistical difference; letters denote significance with a posthoc Tukey test. Alpha = 0.05.

**S2 Fig.**
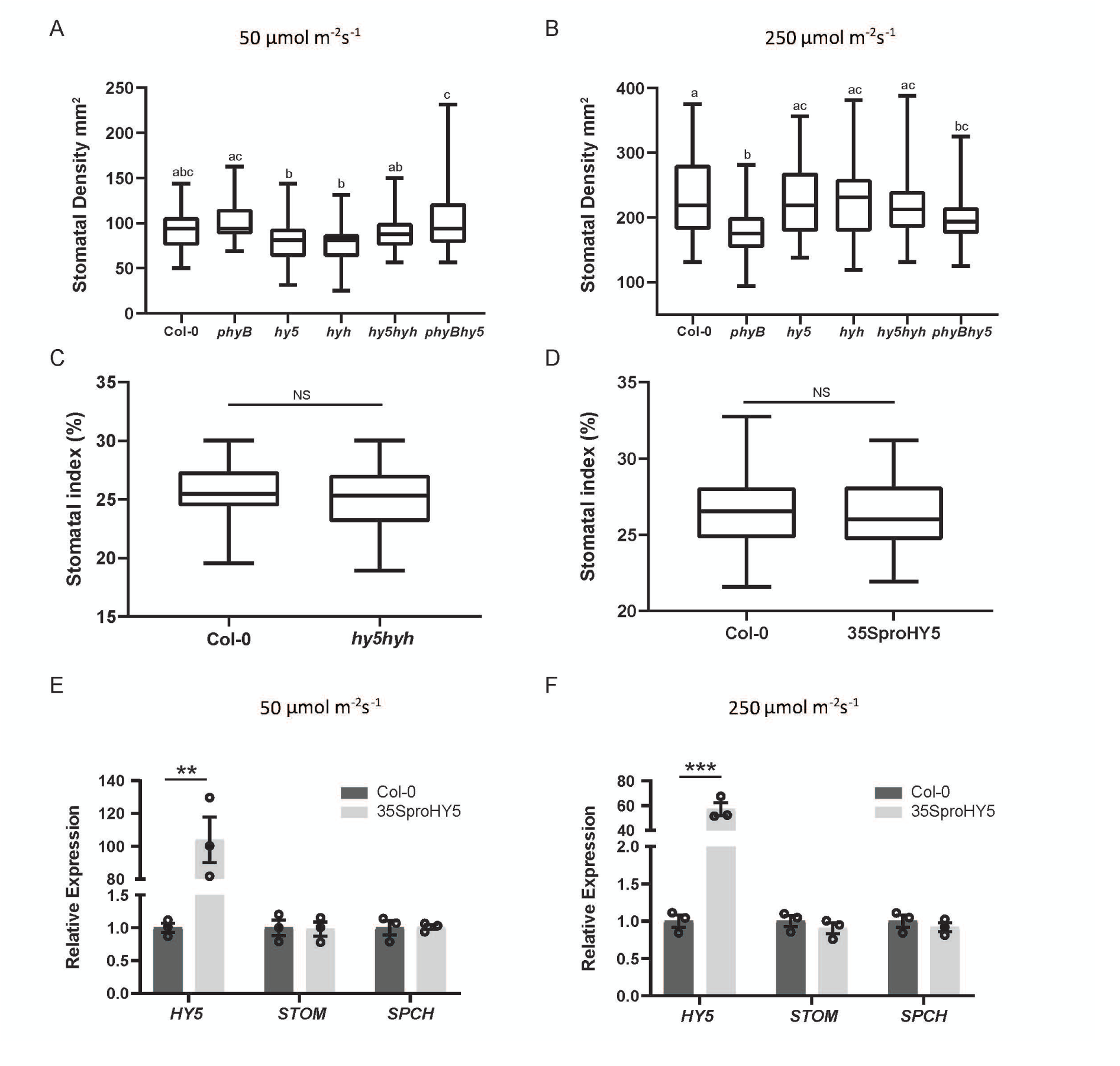
*HY5* is not involved in light regulation of stomatal development. (A) Stomatal density (mm^2^) of Col-0, *phyB*, *hy5*, *hyh*, *hy5hyh*, and *hy5phyB* grown under low light (50 µmol m^−2^ s^−1^). Box plots indicate the 25^th^ and 75^th^ quartiles with the line representing the median; whiskers are the minimum and maximum range. One-way ANOVA was performed to test statistical difference; letters denote significance with a posthoc Tukey test. Alpha = 0.05. (B) Stomatal density (mm^2^) of Col-0, *phyB*, *hy5*, *hyh*, *hy5hyh*, and *hy5phyB* grown under high light (250 µmol m^−2^ s^−1^). Box plots indicate the 25^th^ and 75^th^ quartiles with the line representing the median; whiskers are the minimum and maximum range. One-way ANOVA was performed to test statistical difference; letters denote significance with a posthoc Tukey test. Alpha = 0.05. (C) Stomatal index of Col-0 and *hy5hyh* cotyledons grown under high light (250 µmol m^−2^ s^−1^). Box plots indicate the 25^th^ and 75^th^ quartiles with the line representing the median; whiskers are the minimum and maximum range. Welch’s t-test was used to test for significance; No significance P= 0.4920 (D) Stomatal index of Col-0 and *35Spro::HY5* mature leaves grown under high light (250 µmol m^−2^ s^−1^). Box plots indicate the 25^th^ and 75^th^ quartiles with the line representing the median; whiskers are the minimum and maximum range. Welch’s t-test was used to test for significance; No significance P= 0.3013 (E) Relative expression levels of *HY5*, *STOM*, *SPCH* in Col-0 and *35Spro::HY5* backgrounds grown under low light (50 μmol m^−2^ s^−1^) examined by qRT-PCR, *UBC21* served as internal control. Experiments were performed in biological triplicate and error bars indicate SEM. Asterisks indicate significant differences between low light and low to high light transfers. (Multiple t-test, with statistical significance determined using the Holm-Sidak method, with alpha = 0.05: **P < 0.01). (F) Relative expression levels of *HY5*, *STOM*, *SPCH* in Col-0 and *35Spro::HY5* backgrounds grown under high light (250 μmol m^−2^ s^−1^) examined by qRT-PCR, *UBC21* served as internal control. Experiments were performed in biological triplicate and error bars indicate SEM. Asterisks indicate significant differences between low light and low to high light transfers. (Multiple t-test, with statistical significance determined using the Holm-Sidak method, with alpha = 0.05: ***P < 0.001).

**S1 Table.**
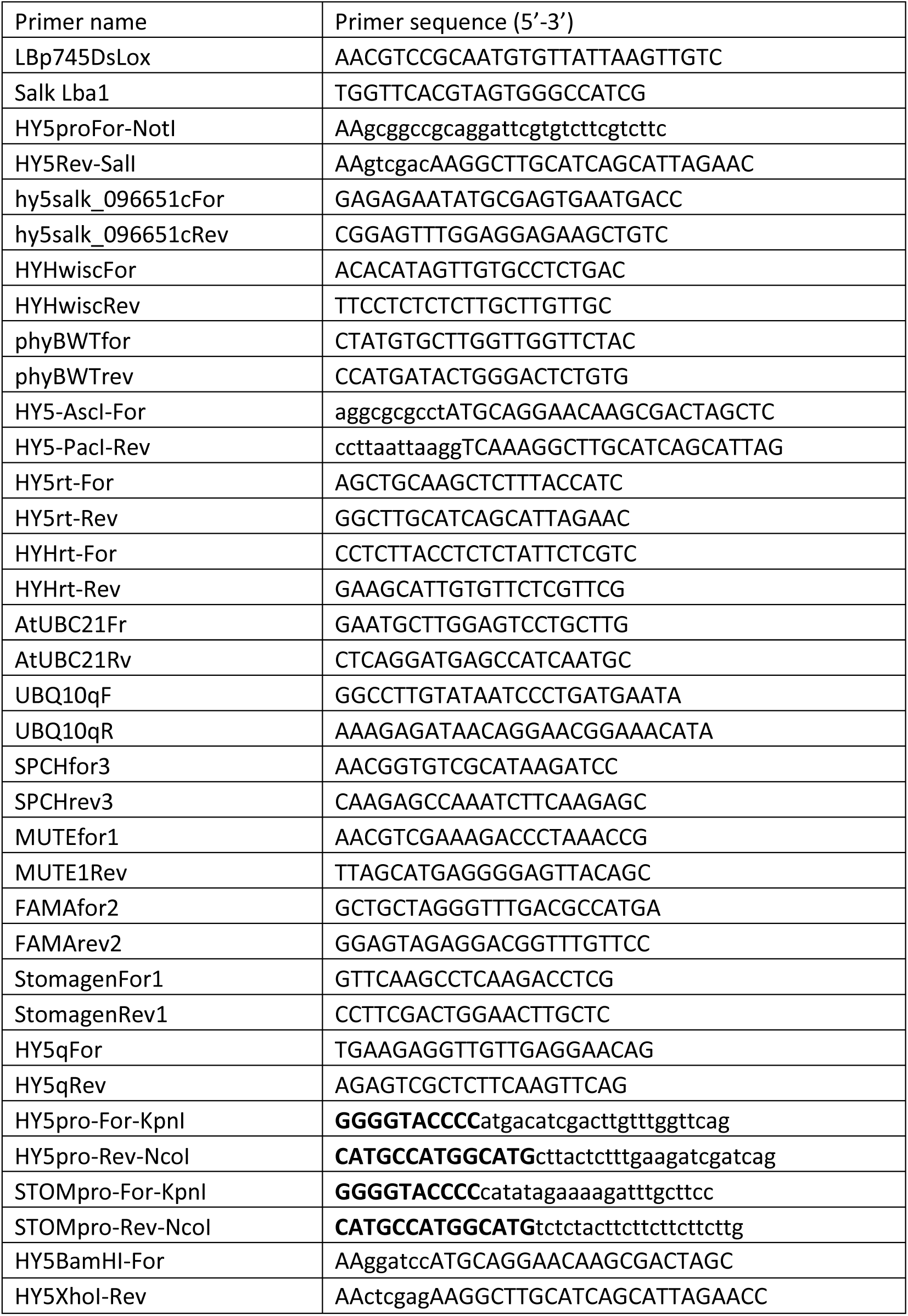
Primer sequences.

